# Data Hunters: bridging bioinformatics education and microbiome research

**DOI:** 10.1101/2024.09.27.615386

**Authors:** Sara Fumagalli, Alice Armanni, Maurizio Casiraghi, Luca Corneo, Giulia Ghisleni, Massimo Labra, Antonia Bruno

## Abstract

Vast sequencing data in public repositories offer researchers unprecedented opportunities for gaining insights through reanalysis and machine learning approaches, thereby advancing the microbiology field. However, the lack of standardized metadata poses a significant obstacle to big data reuse, necessitating advanced bioinformatics skills for curation and integration.

To tackle this challenge, we organized the Data Hunters Workshop, enabling extensive human skin microbiome metadata curation through student community engagement. The Data Hunters Workshop constitutes a student-science initiative within the University of Milano-Bicocca, Italy. Integrating theoretical and hands-on phases, the workshop’s structure included a 6-hour lecture, covering metagenomics and metadata manipulation fundamentals, followed by an interactive practical phase designed as a learn-and-play activity. This phase was supported by in-house developed educational online and command-line resources, facilitating students’ acquisition of Python commands for managing microbiome metadata. Upon the workshop’s conclusion, its effectiveness was assessed through surveys and standardized evaluation scales.

Twenty-nine students addressed the critical challenge of metagenomics metadata standardization. Workshop’s resources have equipped students with foundational knowledge in metagenomics and Python programming, empowering them to curate metadata from 379 amplicon-based and shotgun sequencing projects of the human skin microbiome. The collaborative endeavors of these students will ensure the construction of a curated metadata collection.

The Data Hunters Workshop established a reusable model for similar educational aims, facilitating metagenomics and bioinformatics teaching and metadata curation. It bridged education and microbiome research by focusing on human skin microbiome metadata curation, ultimately striving to construct a specific collection promptly accessible to researchers.

## Introduction

Biology research is undergoing a significant shift as the discovery process has increasingly incorporated quantitative and data-driven approaches alongside traditional hypothesis-driven methods (Işık et al., 2023). Technological advances and the associated cost reduction have led to more accessible large-scale sequencing, resulting in vast amounts of data now available in public repositories. These datasets have proven invaluable across various biological fields, particularly in metagenomics, where the complex nature of the microbiome demands the integration of multiple datasets to extract meaningful insights. Consequently, the demand for highly specialized researchers capable of analyzing and interpreting these large datasets has grown substantially in the last years (Melendrez et al., 2021), and notably, the global bioinformatics market is projected to grow further, reaching $18.96 billion by 2026 (Data, 2019).

In response to the growing demand from both academia and industry for biologists with interdisciplinary profiles that integrate biology and bioinformatics (Attwood et al., 2019; Pucker et al., 2019), many wet-lab researchers have begun, and many others continue to, embark on a bioinformatics journey out of necessity, spending less time to bench work for which they were originally trained (Merali, 2010; Preeyanon et al., 2014). However, engaging with computational workloads that rely on mathematical and informatics methodologies can be challenging without a solid foundation in bioinformatics (Brandies & Hogg, 2021) and prone to errors in results.

The imperative to educate biologists in bioinformatics has reached a critical juncture, making it both essential and urgent. Efforts to address this issue have been ongoing for the past 25 years, from calls to narrow the “skills gap” (MacLean & Miles, 1999) and incorporate mathematics and quantitative thinking in science curricula (Bialek & Botstein, 2004) to the bloom of bioinformatics courses as noted by Tan et colleagues (Tan et al., 2009). Despite this, bioinformatics training continues to be acknowledged as the most unmet need by life scientists (Attwood et al., 2019; Williams et al., 2023). By reviewing higher education institutions’ curricula, it emerged that the progress in integrating bioinformatics fundamentals into biologists’ education remains too slow to ensure that students acquire the necessary bioinformatics skills and competencies before their degree (Attwood et al., 2019; Işık et al., 2023; Melendrez et al., 2021; Williams et al., 2019). Indeed, this training effort must be extended to all biology students, as even a basic bioinformatics education is now essential also to students who do not aim at becoming bioinformaticians (Attwood et al., 2019). Such knowledge enables them to maximize the use of their own data, communicate effectively with bioinformatics colleagues, and gain a broader understanding of the research process (Attwood et al., 2019; “Spotlight on Bioinformatics,” 2016; Tan et al., 2009).

Moreover, the urge to integrate bioinformatics education into biological degree programs is supported by the evidence that bioinformatics training should be introduced early at the undergraduate level (Melendrez et al., 2021; Welch et al., 2014). This approach would help properly address a multidisciplinary field while radicating the essential skills required (Anton Feenstra et al., 2018; Attwood et al., 2019; Rosen & Hammrich, 2020). Without this background knowledge, perceived skills gaps may persist and postgraduate education alone may not be able to address them (Gao & Guo, 2023; Hack & Kendall, 2005), representing a major impediment to research advancement (Attwood et al., 2019).

In recent years, course-based undergraduate research experience (CURE) courses in metagenomics have represented a modern and effective solution to address bioinformatics education challenges in microbiome fields. By combining learning with practical application, CURE courses provide students with essential tools and opportunities to apply them, surpassing the traditional lecture format approach that could limit a strongly applicated field such as bioinformatics (Auchincloss et al., 2014; Gao & Guo, 2023; Lentz et al., 2017). Alongside CURE courses, an international bioinformatics education community strives to enhance bioinformatics training. This community, through collaborative efforts, is establishing guidelines and resources to help instructors’ training and bioinformatics education worldwide spread (Işık et al., 2023).

Driven by the urge to address the pressing challenges in bioinformatics education and contribute alongside the CURE courses and the international community’s efforts, we organized the Data Hunters workshop at the University of Milano-Bicocca, Italy. By bridging bioinformatics training with real-world microbiome applications, Data Hunters directly engaged students in scientific research to face metadata standardization, a central issue in the metagenomics field. Descriptive information of data (Gray et al., 2005), which provides a comprehensive context for samples, metadata constitute the foundational resource for performing analyses and re-analyses (Kumar et al., 2024). This is especially true in metagenomics due to the microbiome context-dependency, resulting in an absolute necessity for host-specific metadata to characterize a microbial community (Knight et al., 2012). However, despite their critical importance, metadata often lack standardization, leading to heterogeneous ontologies use and potential significant information missing that hinder data harmonization and integration (Kumar et al., 2024). Additionally, metadata often include errors, which appear to be unavoidable, even in self-reported data or well-funded clinical studies (Archer et al., 2015; Debelius et al., 2016). As a result, a wide range of curated databases and collections of metagenomics data and metadata have been developed in the last years (Sengupta et al., 2023). In this regard, we initially developed the SKIOME collection, a comprehensive repository of human skin microbiome metadata (Agostinetto et al., 2022). In the last months, we have focused on updating and enhancing this collection by retrieving new metadata from amplicon-based and shotgun sequencing projects, in alignment with FAIR (Findable, Accessible, Interoperable, Reusable) principles (Wilkinson et al., 2016).

The SKIOME collection provided an ideal platform for teaching Python to students and metadata management, offering them a hands-on immersion into the field of metagenomics and bioinformatics. Leveraging SKIOME’s resources, the Data Hunters workshop represents a novel crowdsourcing initiative where students actively participate in curating metadata as an achievement of the skills they acquired through an interdisciplinary lecture and a bioinformatics learn-and-play training. Here, we present the structure and outcomes of the Data Hunters workshop as a potential model for advancing bioinformatics education and microbiome research training.

## Material and methods

The Data Hunters workshop was structured as a four-phase model designed to optimize both learning and practical application. The phases consisted of: (i) an initial in-person lecture, (ii) autonomous learning, (iii) an operative session, (iv) an assessment of the workshop’s effectiveness through surveys and evaluation scales. Each phase is described in detail in the following section.

### In-person lecture

The Data Hunters workshop kicked off on February 28, 2024, with an initial in-person lecture at the University of Milano-Bicocca, Italy. A total of 29 students from the Department of Biotechnology and Biosciences, and the Department of Informatics, Systems and Communication of the University of Milano-Bicocca attended the lecture. The lecture was held in a computer laboratory to ensure that each student had access to a computer.

Reflecting the dual components of the workshop, the 6-hour lecture was structured to equip students with fundamental aspects of metagenomics and bioinformatics equally. The metagenomics lesson was delivered by our team’s wet lab microbiologists to ensure high levels of accuracy and expertise. During the 3-hour lecture, we provided a broad overview of metagenomics, covering from background aspects to details of wet lab techniques to prepare students with the necessary biological knowledge for the operative session of metadata curation. We also introduced the nine categories of information (hereafter referred to as items) that students would identify and verify during the metadata curation process. We dedicated the last part of the lecture to a practical exercise to put in action the newly learned theory. Thus, students were asked to search the nine items from two selected scientific publications about human skin microbiome reproducing the initial process of the manual curation phase. After completing the exercise, a collective discussion was held to verify the results collaboratively.

In the afternoon, a bioinformatician from our team, alongside a researcher in informatics, held the 3-hour bioinformatics lecture. The lecture began with a brief introduction about metadata, public databases, and typical bioinformatics environments such as the terminal and virtual machines. The lecture was organized to alternate between theory and practice, during which students were guided in accessing and navigating a dedicated virtual machine. In the final part of the lecture, the students were introduced to the programming language of Python and they were challenged to solve some exercises.

Since the workshop’s activities following the initial lesson would be carried out remotely, we created a virtual classroom using the app Google Classroom (classroom.google.com) to share announcements, distribute educational materials, and maintain a sense of community among the students.

### Autonomous learning

After the in-person lecture, an autonomous learning phase began, during which students independently gained skills in Python programming and metadata manipulation at their own pace. To support and guide students during the learning phase, we developed two educational resources. The first resource consisted of a GitHub page (https://biomeresearchteam.github.io/data-hunters-chapters/) designed to cover the fundamental aspects of Python, providing students with the necessary commands to manipulate metadata. The resource was structured as theoretical chapters, each followed by exercises specifically designed to reinforce the newly introduced concepts and practice the commands necessary for manual curation. Exercises were then followed by the corresponding solutions and explanations. The second resource was a command-line tool (https://github.com/BiomeResearchTeam/data-hunters-cl-resource) that retraced the steps of our protocol for metadata curation, allowing students to associate each step with the corresponding command. This resource was designed with three levels of Python programming difficulty, enabling students to select the level that suited their skills. If they encountered difficulties, hints and solutions were available for each step. Students run the command-line resource on Deepnote (https://deepnote.com/).

All the resources provided to students were written in Italian to enhance clarity and facilitate understanding.

### Operative session

Metadata from 379 amplicon-based and shotgun sequencing projects related to the human skin microbiome were distributed among the 29 students attending the in-person lecture. Students received a personalized folder containing their assigned metadata files in TSV format, with one metadata file for each microbiome project. They curated metadata files using Python in the Deepnote notebook, following the protocol outlined in the command-line resource.

For each project, students started accessing the corresponding publication by selecting the DOI (Digital Object Identifier) included in the enriched metadata files. These publications provided essential information about samples and methodologies used to generate the sequencing data stored in public repositories. By reading the publication, students determined whether the project concerned human skin microbiome; if it was not, they excluded the project from curation, as it would not be included in SKIOME collection. Conversely, if a project was related to the human skin microbiome, students searched for the nine items in the publication, eventually including supplementary material and cited articles. They then cross-checked these values against the metadata file to update and enrich it in case of any missing or incorrect information.

During the metadata curation process, we offered continuous tutoring support to assist students with both “biological” challenges, such as retrieving information from publications, and “bioinformatics” issues, such as understanding Python concepts, correcting command syntax, and selecting the most appropriate commands for specific cases.

At the end of the metadata curation process, students compiled a detailed report indicating the curated items for each project, any additional information they considered relevant, and, eventually, reasons for excluding projects.

Students who completed at least 75% of the assigned projects received a certificate of participation and academic credit recognition. Additionally, they were acknowledged as co-authors in the resulting publication from the workshop.

### Workshop’s effectiveness evaluation

After the conclusion of the Data Hunters workshop on May 15, 2024, we assessed its effectiveness through a survey consisting of three distinct sections.

In the first section, we utilized a customized Skills of Science Inquiry (SSI) scale (Phillips et al., 2017) to evaluate the perceived skills gained by students during the Data Hunters workshop. This scale comprised eight self-report items, with responses ranging from “Strongly Disagree” (1) to “Strongly Agree” (5), focused on aspects of metadata management and curation. Following the authors’ recommendations, the SSI scale was administered retrospectively to evaluate both the students’ perceived skills after and before attending the workshop for a total of sixteen self-report items.

Students’ confidence in learning and performing metadata curation was measured with a customized Self-Efficacy for Learning and Doing Science (SELDS) scale (Porticella et al., 2017). The scale consisted of eight sentences: four focused on learning and four on performing tasks related to metadata concepts and curation. Students were asked to evaluate each statement using a rating scale from “Strongly Disagree” (1) to “Strongly Agree” (5).

In the last section, we evaluated students’ understanding of the scientific process using both an open-ended question and a true-false test, following the approach outlined by (Brossard et al., 2005). For the open-ended question, students were first asked to self-assess their understanding of the term “human microbiome metadata curation”. If they rated their understanding as “clear” or “general”, then they were asked to provide a description of what they thought the term meant. For the true-false section, students answered twelve statements equally divided between metagenomics and bioinformatics topics covered during the in-person lecture and the autonomous learning phase. Each correct response earned 1 point, while incorrect responses earned 0 points, resulting in a total possible score ranging from 0 (no knowledge) to 12 (high knowledge).

Additionally, we contacted students who did not complete at least 75% of the workshop activities to understand the main reason behind their inability to finish. A brief survey was administered for this purpose.

## Results

A total of 29 students participated in the Data Hunters workshop, engaging in the curation of metadata from 379 amplicon-based and shotgun sequencing projects, with each student responsible for 13 or 14 projects. At the conclusion of the workshop, 17 students achieved the curation of at least 75% of their assigned projects, resulting in the successful curation of 245 projects.

From the post-workshop questionnaire, we received 12 responses, all from students enrolled in the Department of Biotechnology and Biosciences of the University of Milano-Bicocca, Italy, who had completed at least 75% of the Data Hunters activities. Due to the small participant size, statistical analyses were not feasible; however, survey responses provided valuable insights into general trends.

### Perceived skills in science inquiry

A customized SSI scale was employed to assess students’ perceived skills before and after their participation in the workshop. A comparison of the average scores across the eight pre-workshop survey items and the eight post-workshop items revealed a notable improvement in perceived skills for all students except two, who reported a decrease in perceived skills or maintained the same level of perception (Fig. 1). The median score for perceived skills before Data Hunters was 1.88 out of a maximum of 5, while for the post-workshop responses was 3.81, demonstrating a clear enhancement in the students’ self-assessed abilities.

**Figure 1.**
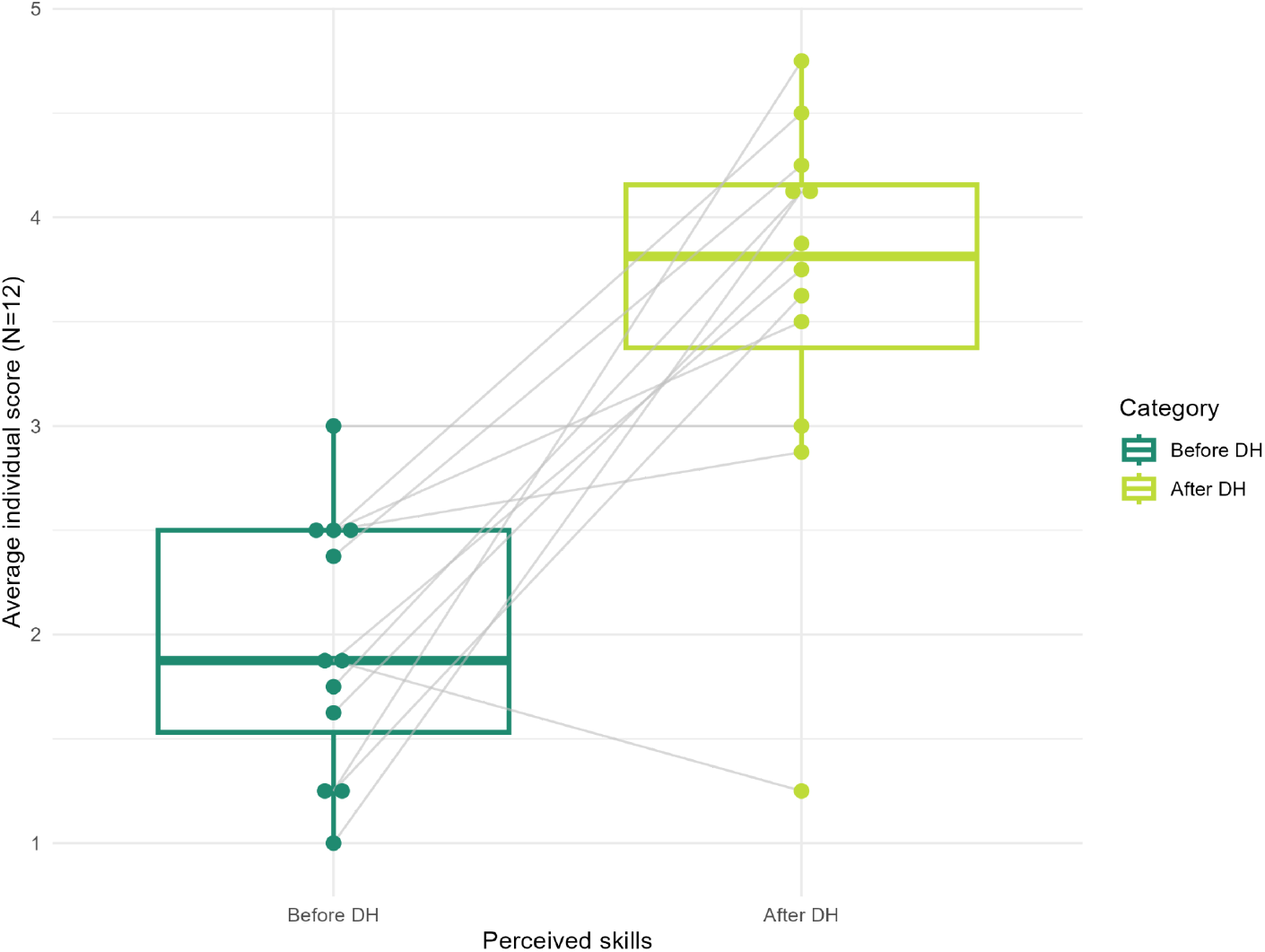
Perceived skills of students before and after participating in the Data Hunters workshop, measured using a customized SSI scale. Points represented the mean scores for eight questions related to before and after the workshop. Gray lines connected the points corresponding to the same student.

### Confidence in learning and doing scientific topics and activities

To evaluate the students’ confidence in learning and performing metadata curation, a customized SELDS scale was applied. A comparison of the average scores across the four learning-related items and the four doing-related items showed comparable results in students’ confidence levels (Fig. 2). Notably, the median scores for students’ confidence in both learning and executing metadata curation were 3.88 out of a maximum of 5.

**Figure 2.**
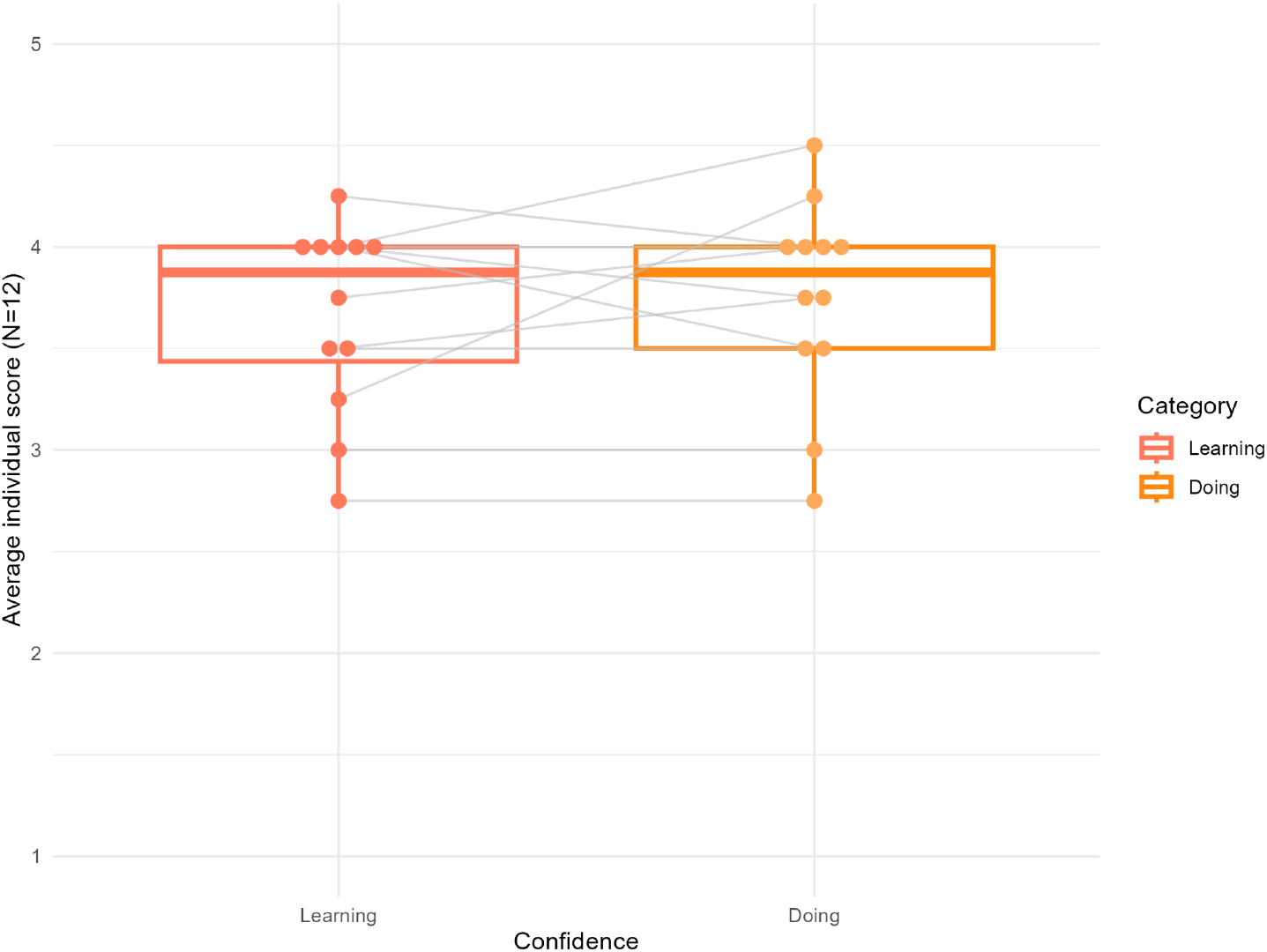
Confidence level of students in learning and doing metadata curation topics and tasks, measured using a customized SELDS scale. Points represented the mean scores for four questions related to learning metadata curation topics and doing metadata curation tasks. Gray lines connected the points corresponding to the same student.

### Understanding of the scientific process

Twelve true-false statements addressing metagenomics and bioinformatics aspects served to evaluate students’ understanding of the scientific process. Students answered between 6 and 11 questions correctly, achieving at least 50% accuracy in their responses. Specifically, four students answered correctly between 50% and 75% of the questions, while eight students scored between 75% and 100%.

Considering metagenomics and bioinformatics questions separately, students exhibited a balanced distribution of correct answers across questions for these two topics, with a slightly higher accuracy in metagenomics (Fig 3).

**Figure 3.**
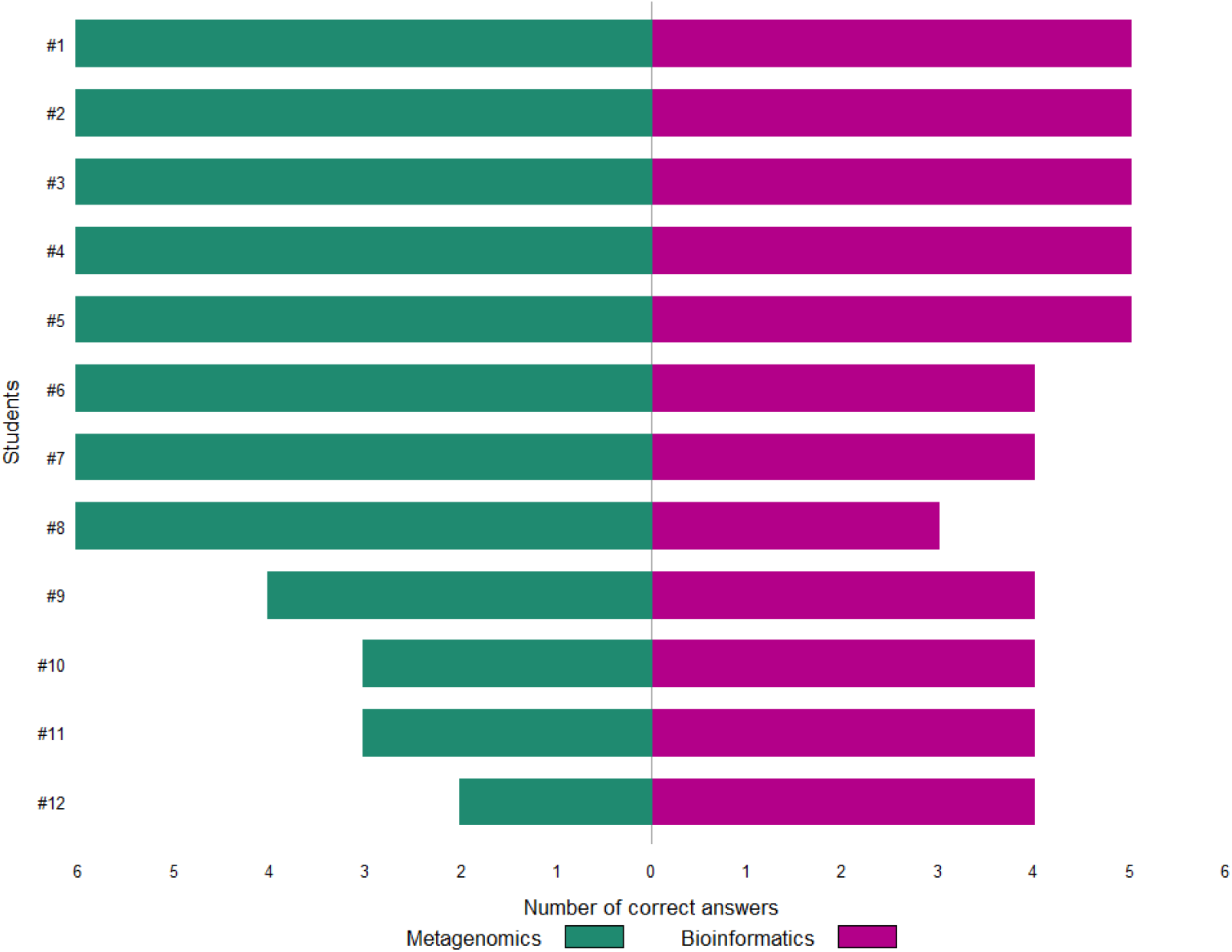
Distribution of correct answers in the true-false test across metagenomics and bioinformatics topics. For each student, the number of correct responses in the true-false test is presented. The maximum number of correct answers for both the metagenomics and bioinformatics topics was 6, resulting in a total maximum score of 12.

We further evaluated students’ perceived skills after the Data Hunters workshop and their confidence in learning and doing, and compared these with their scores on the True-False test. Our findings indicated that 75% of the students (N=9) underestimated their abilities, while 17% (N=2) overestimated their skill level. 8% (N=1) accurately assessed the skill level. (Fig. 4).

**Fig 4.**
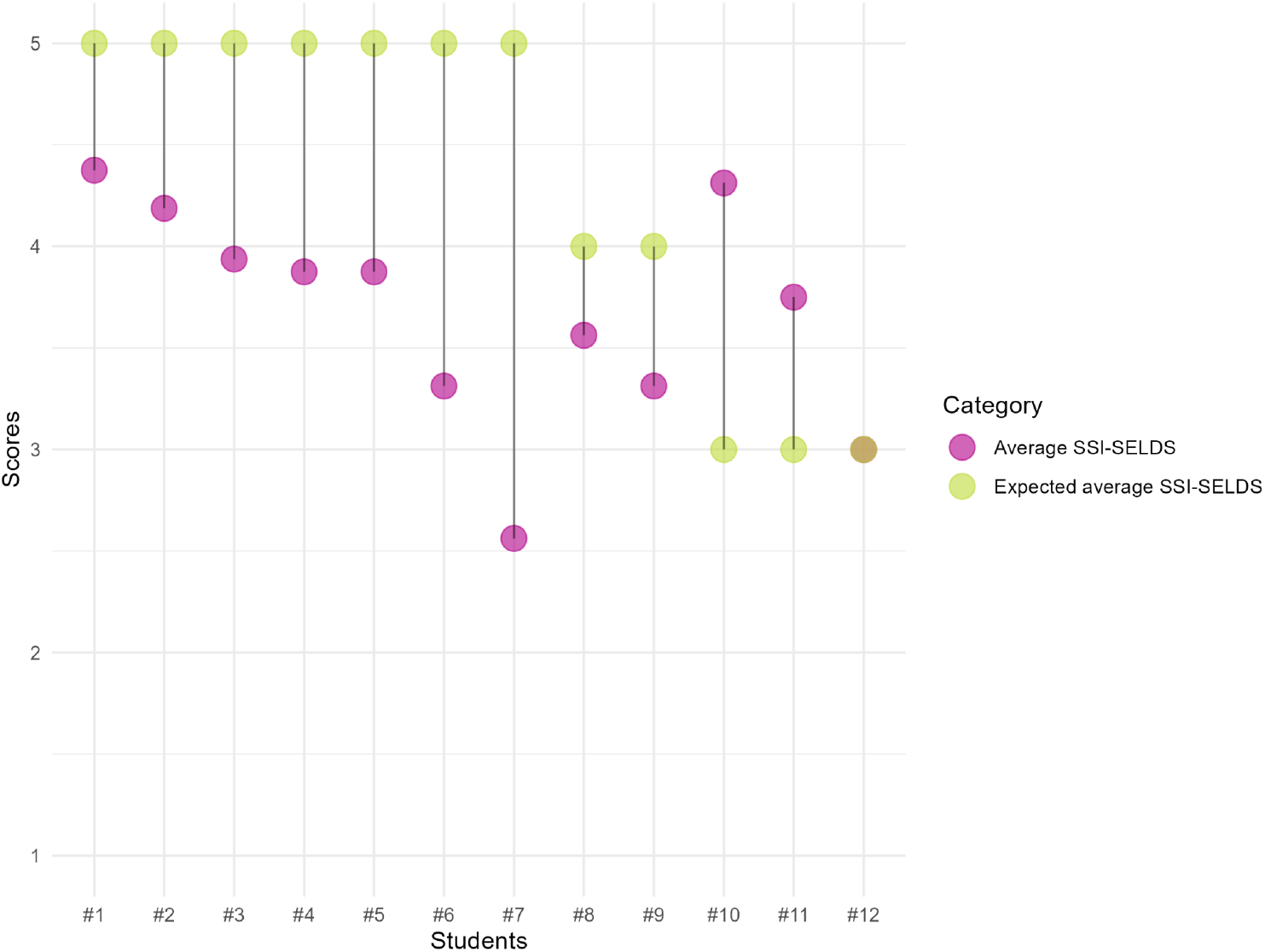
Average value of SSI after Data Hunters and SELDS compared to expected values derived from the true-false test results. For each student, the average of the post-workshop SSI and SELDS scores is presented in purple, while the total score from the true-false test converted to a 1-5 range is shown in green.

Lastly, we received 10 responses from students who did not complete at least 75% of the activities of Data Hunters. Eight of them stated that the workload was incompatible with their other commitments, such as study and exams, while two students responded that the workshop did not meet their expectations.

## Discussion

The Data Hunters Workshop represents a novelty in bioinformatics education, particularly in the field of microbiome research. This workshop was the first of its kind to directly address the critical issue of human skin microbiome metadata curation through a structured and hands-on educational approach. By focusing on the integration of education and real-world bioinformatics context, the workshop set a precedent for integrating metagenomics teaching with actionable research tasks. The hands-on curation of 245 human skin microbiome projects, in addition to providing an invaluable resource, gave students direct experience in addressing one of the most significant barriers to the reuse of public microbiome data, namely the lack of standardized metadata. The workshop’s success in equipping students with the skills to overcome this barrier represents a significant advancement in bioinformatics education, with the potential to be adapted and reused in similar settings.

Moreover, the Data Hunters workshop enabled students to address a critical metagenomics challenge through a collaborative endeavor, promoting the development of a robust interdisciplinary background. Designed for university students, Data Hunters adhered to the commitment of introducing scientists to bioinformatics skills early in their academic journey, helping their skill maintenance and proficiency (Attwood et al., 2019).

Data Hunters demonstrated a remarkable positive impact on the acquisition of bioinformatics skills related to metadata curation process, as evidenced by the results in Fig. 1. The considerable increase in perceived competence among students after attending the workshop highlights the effectiveness of the pedagogical model employed. By providing students with opportunities to work on real scientific research issues, a deeper understanding and a sense of ownership over the learning process was achieved. As previously reported (Pucker et al., 2019; Williams-Pierce, 2011), active involvement in experiments promotes a greater sense of motivation in students, as it allows them to experience success that subsequently fosters self-confidence and propensity for learning (Williams-Pierce, 2011). The challenge lies in structuring a balanced educational activity that equips students with the necessary skills to face tasks that must be achievable to ensure a successful experience (Williams-Pierce, 2011) and engaging to keep them motivated. In this regard, our results revealed that students rated their confidence in learning theoretical concepts and performing practical tasks at comparable levels, as shown in Fig. 2. This indicates that the workshop successfully promoted a balanced development of both knowledge and hands-on bioinformatics skills applied to the metagenomics field.

The True-False test provided a valuable assessment of the overall students’ understanding of the metadata curation process, as well as the actual skills they acquired during the workshop. Moreover, by testing both metagenomics and bioinformatics questions, we aimed to assess the efficacy of an integrated approach. The results were considerable, with all students achieving at least 50% correct answers. As illustrated in Fig. 3, even those with lower scores demonstrated a solid understanding of both topics, with no students answering all questions incorrectly. This outcome is particularly noteworthy given that the students who filled the survey were all from the Department of Biotechnology and Biosciences, where no dedicated course in metagenomics or bioinformatics exists. However, while students are expected to have a foundational biological competence to engage with new biological topics, there are no similar expectations for bioinformatics. Notably, the slightly higher performance in metagenomics may be attributed to students’ stronger familiarity with biological concepts, whereas bioinformatics represents a new domain of learning for them.

We can attribute these positive outcomes to the educational model we developed, which alternates traditional lectures with autonomous learning phases. By granting students greater independence during practical activity, we can enhance their interest and self-confidence. When students take responsibility for their learning, they demonstrate higher levels of commitment and self-esteem (Williams-Pierce, 2011). Additionally, it is crucial to customize the learning process to align with each student’s unique needs and goals. For this reason, the Data Hunters workshop included an autonomous learning phase that allowed students to train at their own pace. Moreover, we implemented a game-based training, through the use of educational learn-and-play resources, to overcome initial barriers to acceptance of new concepts, and promote students’ motivation and engagement, as suggested by other works (Venkatesh, 1999; Williams-Pierce, 2011).

However, as illustrated in Figure 4, it is evident that most students underestimated their abilities relative to their performance on the true-false tests. While this discrepancy may be influenced by the simplicity of the true-false questions, which might not fully capture the depth of the learning and working process, we believe it serves as a valuable tool for assessing students’ self-evaluation capabilities and workshop’s efficacy. The tendency for students to underestimate their skills may derive from the limited tools available to fully recognize their competencies, particularly due to the novelty of bioinformatics topics. This unfamiliarity with the field could impact their ability to appreciate their complete range of skills. This observation highlights a future opportunity to integrate some forms of feedback into our educational model, enabling students to develop the resources necessary for more accurate self-assessment.

In conclusion, the Data Hunters workshop has proven to be an effective educational model for metadata curation, gaining significant results in enhancing students’ bioinformatics skills. By bridging education with microbiome research, the Data Hunters workshop constitutes a scalable model that can be adapted for diverse educational objectives, exceeding our focus on human skin microbiome metadata. This initiative not only addresses the existing skills gaps in bioinformatics among biologists but also contributes to the standardization of metadata practices. Moreover, as a concrete effect of the collaborative efforts of our participating students, we have successfully curated a significant portion of the SKIOME collection, achieving results that would have been unattainable independently and, thus, establishing a vital resource to drive future advancements in microbiome research.

